# Protein-Ligand Docking with Protein-based and Ligand-based Structure Activity Relationships

**DOI:** 10.1101/394874

**Authors:** Darwin Y. Fu, Jens Meiler

## Abstract

Protein-small molecule docking programs predict the interaction interface and energy between a given protein target and a small molecule ligand. The accuracy of docking predictions generally improve with the guidance of experimentally derived restraints. One available source of such restraints is structure-activity relationships (SARs). SARs provide information on changes in binding affinity or biological response corresponding to a chemical change in the protein and/or ligand. These chemical changes frequently refer to amino acid mutations on the protein side and functional group modifications on the ligand side. Theoretically, predicted interaction energies should correlate with SARs though in practice, this is challenging due to the difficulties in scoring protein-ligand interactions. We have previously developed RosettaLigandEnsemble (RLE), a protein-ligand docking method that simultaneously docks a congeneric ligand series to a single protein target. RLE is capable of identifying native-like binding modes for a ligand series that match the available ligand SARs. This work in progress reports on the extension of RLE to factor in SARs derived from protein mutagenesis data. The new method, ProteinLigEnsemble (PLE), is also part of the Rosetta Biomolecular Modeling Suite available at https://www.rosettacommons.org/. We have also developed protein ensemble docking features that allow for docking or screening against multiple receptor variants at the same time. We have included a proof of concept study and a tutorial for interested users.

## Introduction

### Ensemble approaches from the protein perspective

RosettaLigandEnsemble [4] was developed to dock a series of congeneric ligands to a single protein target with the option to include structure-activity relationship (SARs) for ligand modifications. As one might imagine, this can be extended to utilize the SARs associated with protein changes rather than ligand changes. Ligand binding changes upon protein residue mutagenesis is often utilized as a way to localize the interaction site. We have extended docking in Rosetta to process mutation data automatically and to use the SARs to guide modeling.

Existing methods make use of protein ensembles for working with challenging protein targets that are not well represented by a single static structure. This often occurs with protein targets with highly flexible binding sites and/or a significant induced fit effect. In one study, Ellingson et. al. used molecular dynamics snapshots to improve decoy discrimination over docking against a single crystal structure [3]. This strategy can also be used to generate the holo protein conformation when starting with an apo structure [6]. Although Rosetta does not perform molecular dynamics, we have now enabled the ability to use protein ensembles with an ”average-grid” scoring method in RosettaLigand [2] and RosettaLigandEnsemble. This greatly increases the amount of protein flexibility Rosetta can process on top of the backbone and side-chain degrees of freedom typically allowed. One interesting feature is that Rosetta protein ensembles do not have to be comprised of the same protein variant. In other words, Rosetta protein ensembles can be used to dock or screen against multiple mutants simultaneously.

### Combining protein and ligand SARs

The new PLE docking approach is designed for use cases where a ligand panel is tested against several protein mutants. By docking every pairwise combination of proteins and ligands at the same time, PLE adds an additional dimension over previous approaches that handled protein and ligand SARs independently. Figure 1 shows the distinction among the datasets for each of the ensemble docking approaches.

**Figure 1.**
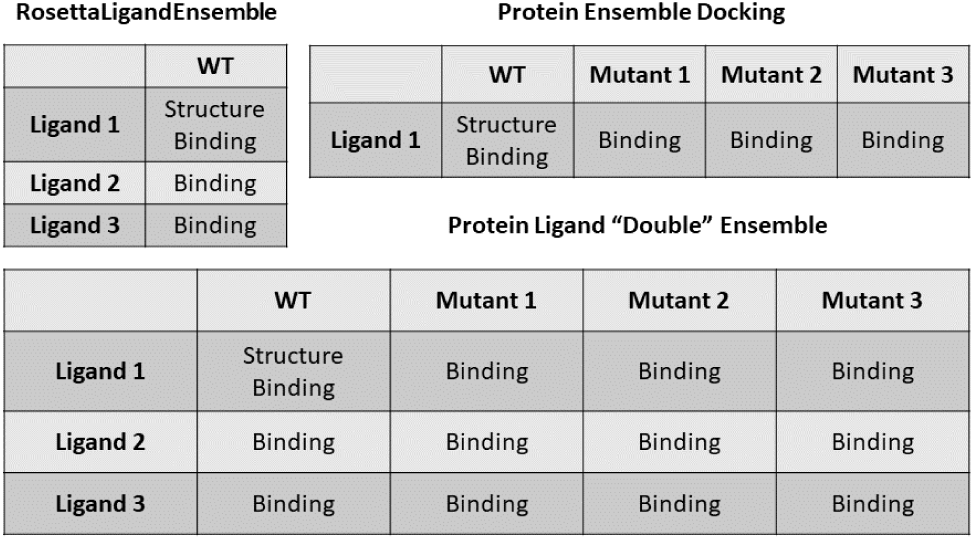
Datasets for various SAR-guided ensemble docking approaches. Experimental datasets that different ensemble approaches are designed to work with. In each case, it is assumed that the ligands have a common central scaffold and that binding data is measured the same way for all protein-ligand pairs. Only a single input structure is required and it does not have to be the wildtype protein.

The PLE method is for the Protein Ligand ”Double” Ensemble docking example. The algorithm will optimize interfaces between wildtype, Mutant 1, Mutant 2, and Mutant 3 Proteins with each of the corresponding Ligands 1, 2, and 3. Essentially, a single PLE docking run on this dataset will generate twelve models. Though the general placement and orientation of the ligands in these models will be similar, the exact interface conformations are simulated independently. An SAR correlation correction factor will promote binding modes where the Rosetta score matches the SAR table. It is important to note that the SAR data table do not have to be complete in order for PLE to be applicable. The algorithm will automatically discount protein-ligand pairs without SAR data from the correlation calculation. In the case where only one ligand is tested against several protein mutants, PLE can also be used to perform docking across the single dimension.

## Materials and Methods

### Ensemble docking workflows

The new ensemble features can be utilized as part of three workflows summarized in Figure 2. Workflow A represents RosettaLigand single protein-single ligand docking protocol described in DeLuca et. al. [2]. The modification now allows alternate receptor conformations to be passed into the low resolution docking phase with the ensemble proteins XML tag. The alternate receptor conformations or mutations are used to calculate the scoring grid in the same manner as the original protocol. A single protein-ligand complex is produced using either the best scoring receptor model or the user’s preferred receptor model.

**Figure 2.**
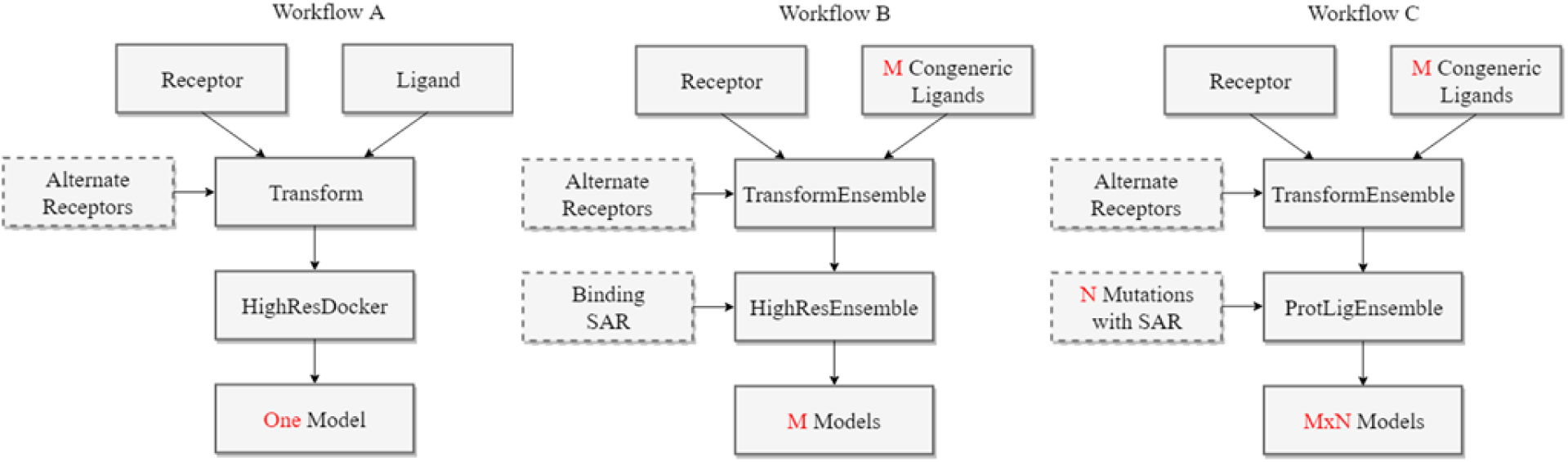
Suggested workflows for ensemble ligand docking approaches in Rosetta. Suggested workflows with optional inputs shown in broken line boxes. **A:** RosettaLigand workflow for using multiple receptor structures. **B:** RosettaLigandEnsemble workflow for using multiple receptor structures and SAR refinement. **C:** ProteinLigEnsemble workflow for double ensemble docking with both protein- and ligand-based SAR.

Workflow B is the multiple ligand-single protein RosettaLigandEnsemble protocol we previously developed. Similarly to workflow A, a new feature allows alternate receptor models to be passed into the low resolution docking phase. A single receptor model is passed on to the high resolution docking phase where SAR guided ligand ensemble docking takes place. Users can specify whether they wish to use the best scoring model or to use the primary input model with the use main model XML tag. The final output contains one protein receptor in complex with each of the input ligands.

Workflow C is the new ProteinLigEnsemble double ensemble method for multiple proteins-multiple ligands docking. The approach in docking is similar to that of RLE in the low resolution phase. Alternate conformations may also be used in this stage. The additional protein mutants are generated in the high resolution PLE stage. Similar to RLE, a rank correlation to experimental data will be used to promote docking modes that generate the most favorable Rosetta scores for the strongest binders. Compounds without binding data to a given receptor will not be factored into the Spearman’s coefficient. The final output contains a structure for every possible pair of input ligands and input proteins.

Additional changes in PLE are the optimization order and the docking radius. The RLE SAR correlation guidance optimized one protein-ligand pair after another in iterative cycles. This has been adjusted to complete all optimization cycles for the protein-ligand pair with the highest experimental affinity first. This should generate the best scoring structures for the strongest binders, improving correlation with experimental data and reducing the stochastic nature of the output model scores. Furthermore, a docking radius has been introduced as a simple way of defining the binding pocket. Previous iterations required individual definitions for each ligand, leading to increasingly bulky XML scripts. The new version automatically defines the flexible portion of the protein structure based on a single setting and applies it to all protein-ligand pairs.

### ProteinLigEnsemble proof of concept dataset

Two test cases are derived from individual studies on two GPCRs, the adenosine A2A receptor and the neuropeptide Y1 receptor. The A2A dataset focus on work by Zhukov et. al. [8] in which a panel of antagonists are tested against point mutants of a thermostablized receptor. Figure 3A shows the binding pocket of the A2A-ZM241385 complex with available mutational data denoted in purple. SCH420814 is a ligand with a significant addition to the extracellular binding end of the ligand. The crystal structure was published separately by Jaakola et. al. (PDB: 3EML) [5]. The Y1 dataset, shown in Figure 3B is derived from Yang et. al., which contains a crystal structure of the Y1 receptor in complex with the antagonist UR-MK299 (PDB: 5ZBQ) [7]. The additional antagonists feature the addition of various linear amide groups to the central scaffold of UR-MK299.

**Figure 3.**
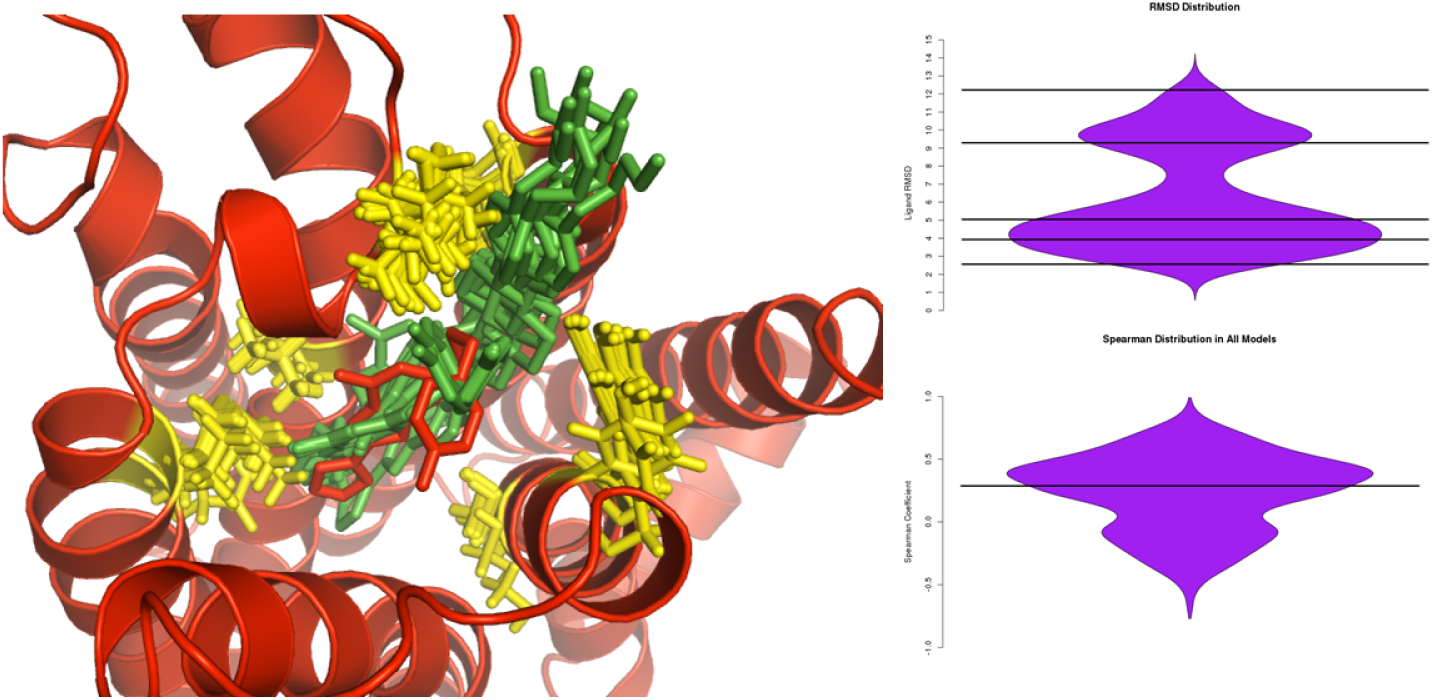
Two proof of concept test cases. The crystal structure is shown in green, the ligand in orange, and the mutation sites are marked out in purple. SAR measurements are given in the tables. **A:** A2A dataset derived from Zhukov et. al. containing binding data (pKd) for two ligands against seven receptor variants. Crystal structure of wild type receptor is shown in complex with ZM241385. **B:** Y1 dataset derived from Yang et. al. containing binding data (nM Kd) for four ligands against five receptor variants. Crystal structure of wild type receptor is shown in complex with UR-MK299 (PDB: 5ZBQ).

## Results

Figure 4 summarizes the docking results for the A2A test case. The docked models capture the general orientation of the binding mode observed in the crystal structure as indicated by the significant portion of docked models around 4 RMSD. However, the compact conformation of the ZM241385 ligand with the wildtype receptor is not well captured. This may be an issue with the pregenerated conformations rather than the docking method. One issue with the RMSD analysis is that only one protein-ligand pair can be compared for native-like binding modes. Alternative measures may be necessary to evaluate the efficacy of the method across the entire dataset. A significant enrichment of binding modes that correlate positively with the SAR is observed for the A2A test.

**Figure 4.**
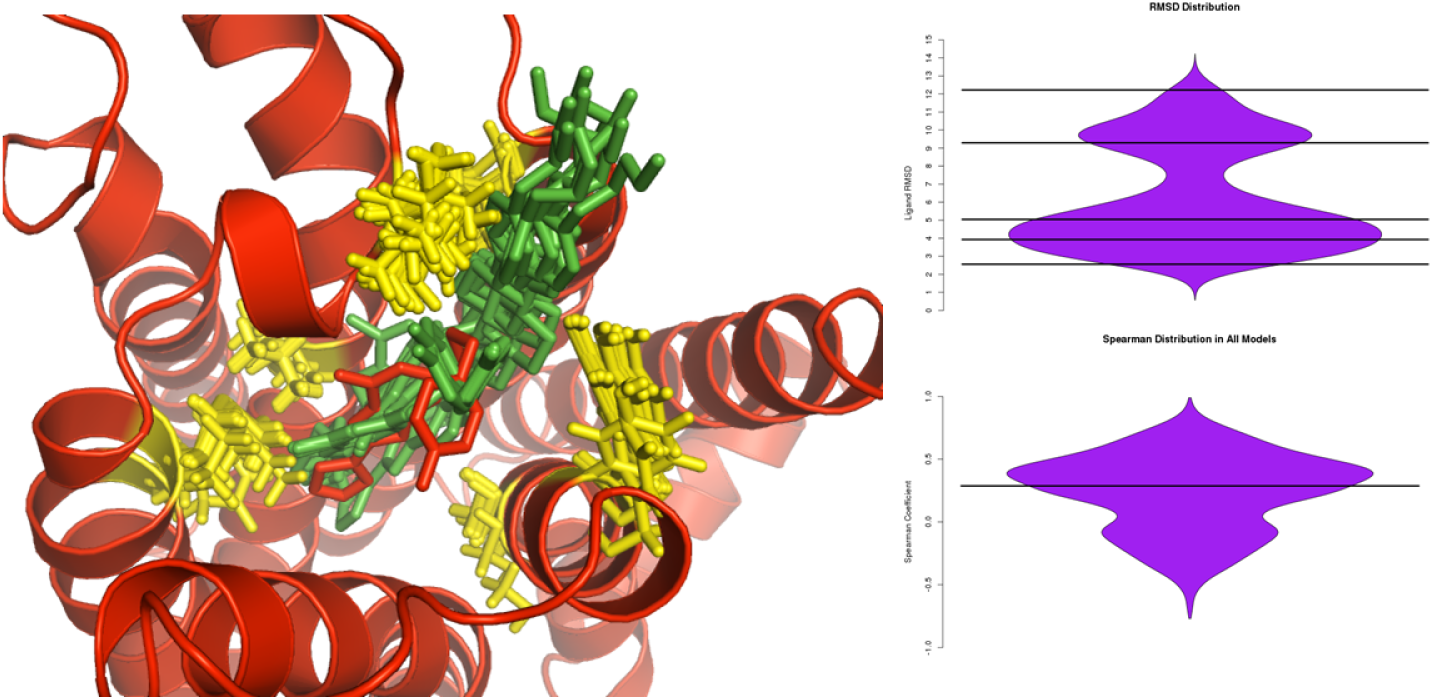
A2A double ensemble docking results. Summary of docking results with top right graph showing the RMSD distribution of wildtype-ZM241385 models, and the bottom right graph showing the experimental rank correlation. A representative output is shown overlaid with the 3EML crystal structure. The crystal receptor structure and bound ligand is in red, docked ligands are in green, and mutation sites are shown in yellow.

Figure 5 shows the docked models from a single simulation in grid format. One benefit of double ensemble docking over individual docking runs is the consistency of docking modes across the entire system. Rather than performing ad-hoc analysis to distinguish putative binding modes generated from individual runs, PLE identifies binding modes consistent with the entire dataset. The interfaces consists of individually optimized ligand conformations and protein side-chain positioning. One limitation of this dataset is that the side-chain mutations consisted of alanine scanning, which in some cases simply represent a reduction in side-chain steric volume. A more extensive dataset including significant electrostatic based mutations could be useful in testing how well PLE captures the SAR of substantial changes.

**Figure 5.**
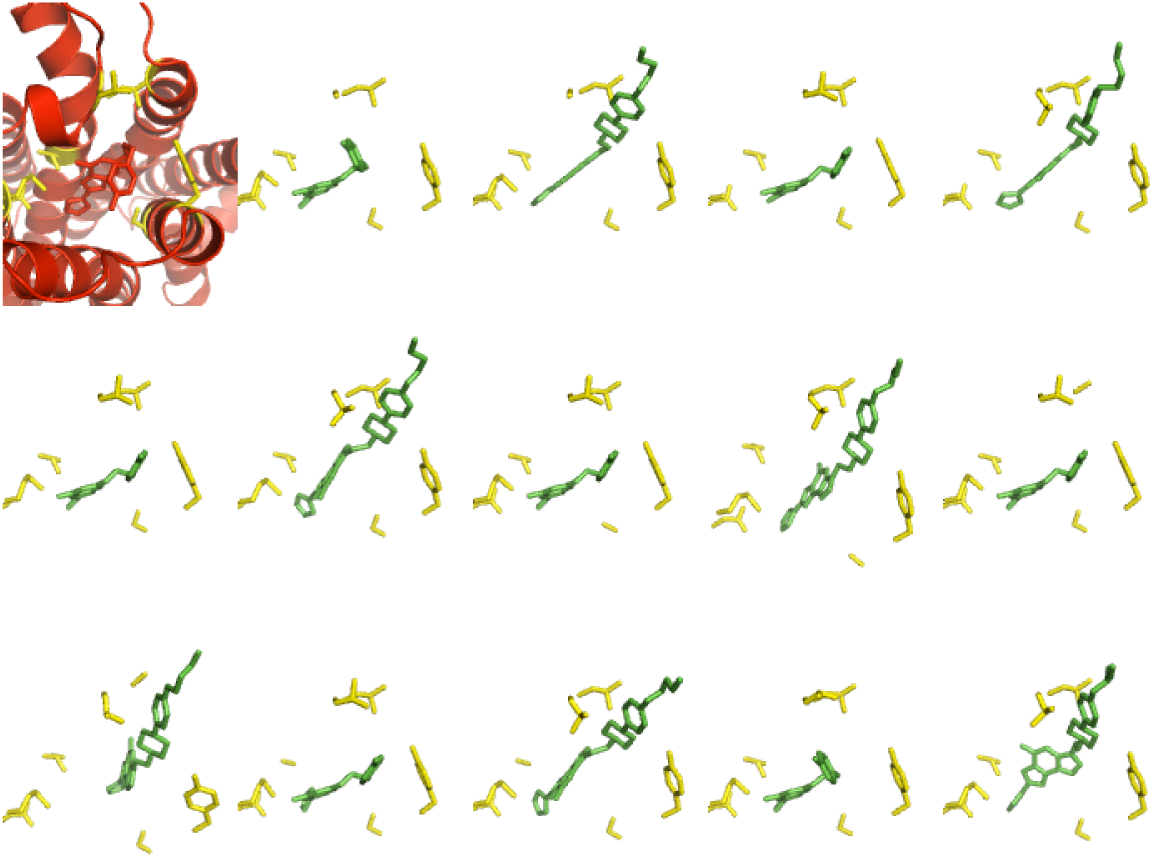
A2A docked ligands in grid view. Docking output from a single run of double ensemble docking. Top left corner shows the crystal structure of wildtype protein in complex with ZM241385. Remaining panels show each of 14 possible combination of 2 ligands with 7 protein variants. The crystal structure is in red, docked ligands are in green, and mutation sites are shown in yellow.

Figure 6 shows the Y1 docking results summary. As indicated by the RMSD distribution, the native like binding mode was captured for a significant number of wildtype-UR-MK299 models generated. One issue that continues to arise is the band of high RMSD models. Like RLE, PLE does not take into account SAR during the low resolution docking phase. Future development could focus on a way to improve model selection with low resolution scoring grids without adding substantial computational cost. The Spearman distribution in the Y1 test case is also less than ideal. This is likely due to the larger number of protein-ligand pairs being modeled. PLE optimizes binding pairs in order of affinity and it is possible that the SAR correlation becomes too restrictive when docking the weaker binding pairs. This is analogous to the difficulties of negative design when using an algorithm designed to find more and more favorable scores.

**Figure 6.**
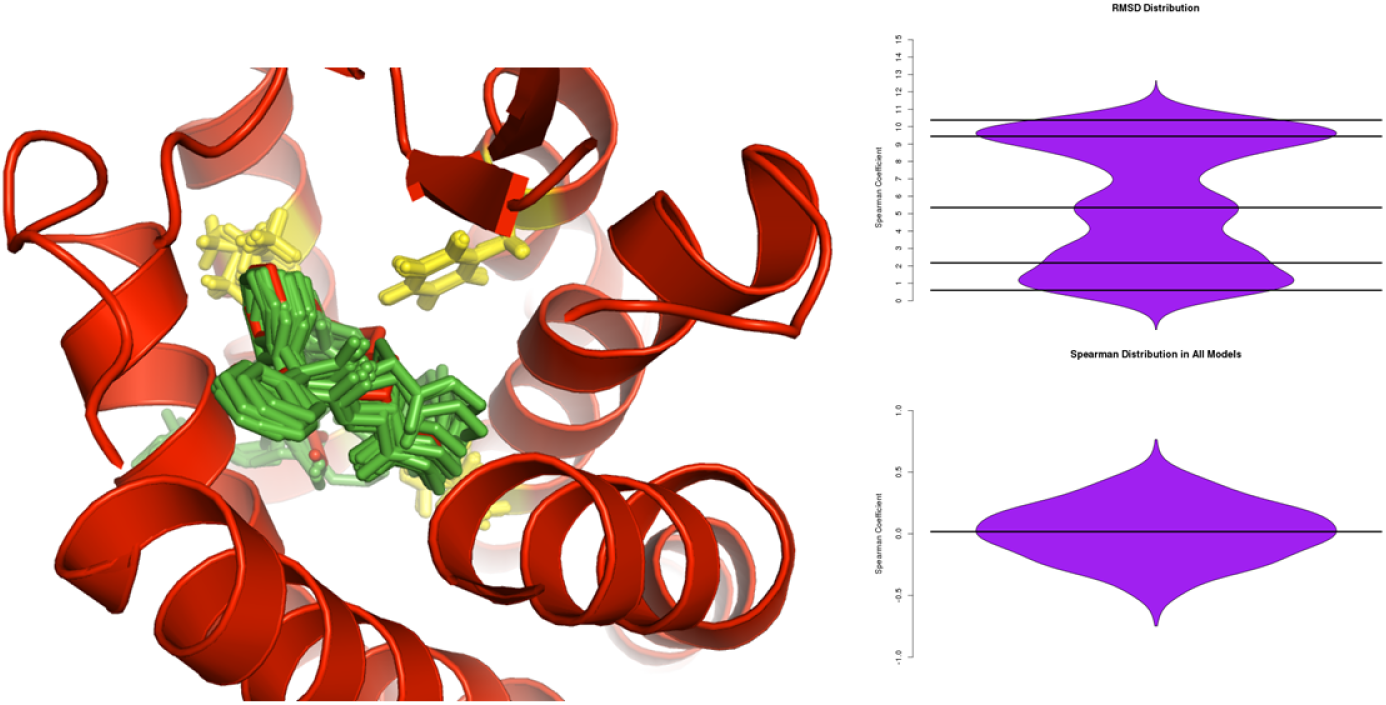
Y1 double ensemble docking results. Summary of docking results with top right graph showing the RMSD distribution of wildtype-UR-MK299 models, and the bottom right graph showing the experimental rank correlation. A representative output is shown overlaid with the 5ZBQ crystal structure. The crystal receptor structure and bound ligand is in red, docked ligands are in green, and mutation sites are shown in yellow.

## Conclusion

Protein ensemble and double ensemble docking (PLE) features have been incorporated into the Rosetta Ligand Docking. The proof of concept shows that PLE is capable of generate a consistent set of docked models with protein-ligand SAR guidance. Additional benchmarking is necessary to explore the full potential and limitations of the new algorithms. One suggestion for benchmarking the addition of protein ensembles to the RosettaLigand and RosettaLigandEnsemble workflows is via PSCDB. PSCDB is a database that tracks different types of receptor motion upon ligand binding [1]. This may be particularly helpful in assessing the use of ensemble methods for ligand binding to highly flexible loop regions.

To augment the double ensemble proof of study benchmark in an application-like way, a GPCR ligands dataset may be a good starting point. Many of these receptor studies include mutational data in their supplemental and there may be additional ligand compounds related to the co-crystallized ligand. It is unlikely a binding value for each protein mutant - ligand pair will be found, but PLE can work around this limitation by ignoring missing SAR values. One other concern for dataset building is that different assays may be used for different protein-ligand combinations. In particular, some of the ligand data may be a measure of biological response, which can be due to allosteric effects as opposed to binding affinity changes. The presence of misfolding mutations is also a concern but these can be experimentally determined without the need for a fully solved structure. Careful curation will be necessary to generate a proper and diverse benchmark dataset.

### ProteinLigEnsemble protocol capture

The following section provides an example for using the double ensemble docking protocol. Rosetta can be obtained through www.rosettacommons.org

All files associated with this protocol capture is provided in the demos/protocol capture /rosettali-gand_ProtLigEnsemble/ directory of the Rosetta distribution. This protocol has been tested to work with Rosetta version 9820fea, released July 12, 2018. Examples commands for this protocol are numbered in the commands file of the protocol capture folder and referenced as (1), (2), (3)… etc.

### Starting files

The raw starting files are a single target protein receptor structure in PDB format, and a series of ligands in SDF format. The receptor structure used in this example is neuropeptide Y1 receptor bound to the ligand UR-MK299 (PDB: 5ZBQ). This file can be found in /inputs/ as protein.pdb. The four congeneric ligands in /prep/aligned ligands/ directory have been aligned by their core scaffold. The reasonable number of ligands depends on the number of protein variants considered as each run generates all possible pairs. We generated up to 30 models per docking run without issue though this number may change depending on your computational setup.

### Ligand preparation

Ligand preparation can be performed in the same fashion as previously documented RosettaLigand docking procedures. Commands 1, 2, and 3 cover the process of generating conformations with the BCL conformer generator and creating Rosetta ligand param files. Any conformation generator can be used as long as it can produce ligand files in the required SDF or MOL format. Example ligand conformers have been created for you in /prep/conformers and the necessary param generation files are provided in /prep/make params/

The final Rosetta input ligand files are provided in /prep/rosetta inputs/. The ligands have been designated with the letters B,C,D,E though you are free to use any chain designation as long as they are different from each other and the protein receptor. The correspondence between published ligand designations and the Rosetta lettering is provided in ligands.list. The params file process is the same as those for RLE but you can skip adding SAR data to the param files as SAR data will be provided in a separate dedicated file.

### Setting up the QSAR file

The QSAR file, an example of which is provided as inputs/qsar.txt, will provide protein-ligand pairs of interest to the ProtLigEnsemble mover. Each line is organized as a protein identifier, a ligand identifier, and an optional binding value. To provide a ligand binding value to a wildtype protein, enter:

WT B 0.17

where WT indicates wildtype, B is the single letter chain of the ligand, and 0.17 is the optional affinity value. Note that the affinity value can be any measure as long as they are self consistent. Rosetta assumes the lower values indicates a more favorable binding. This can be changed by setting a negative correlation weight in the -docking:ligand:ligand ensemble option. To provide a binding value to a mutant protein, enter:

107 A B 7.5

where 107 A indicates a mutation at residue 107 to alanine, B is the single letter chain of the ligand, and 7.5 is the optional affinity value. This will cause Rosetta to generate a 107A mutant regardless of what the wildtype residue at position 107 is. Note that this numbering system must correspond to Rosetta pose numbering, where the first residue is numbered 1, the second is numbered 2…and so on. For the time being, PLE is designed to work with single mutants. This QSAR file will be provided to Rosetta in the XML script as the qsar file tag for the ProtLigEnsemble mover.

### Input file organization

For PLE runs, it is preferred to combine all aligned ligand PDBs into a single PDB file. This is provided as ligands.pdb in the /inputs/ folder. Note that the params and conformers files are not combined, just the single ligand PDB inputs.

In addition to structural files, a RosettaScripts XML file and a Rosetta options file. The XML file describes the custom protocol to be used by Rosetta. Details of how to setup an XML file and the meaning of the individual tags can be found by searching the documentation website https://www.rosettacommons.org/docs/latest/. The example dock.xml provided uses the settings from the benchmark. Actual application use may require the user to alter these values according to biological context. The defined scoring function is based on the existing RosettaLigand scoring function, but may be substituted in the XML script. The provided options file defines Rosetta input and output directories along with a number of sampling parameters. A full options list is available on the documentation website. The ligand ensemble option is necessary to use PLE; a weight of 0 can be used to run PLE without taking SAR data into consideration.

A few XML tags in the ProtLigEnsemble mover are newer features to this mover. The aforementioned qsar file tag tells Rosetta where to find the QSAR file. The distance tag defines the radius, in angstroms, of the sphere around the ligand considered to be the binding pocket. All residues in this sphere are considered to be flexible. This tag replaces the LigandArea, InterfaceBuilder, and MoveMap tags RosettaLigand users may be familiar with. The ignore correlation option tells PLE to avoid calculating the rank correlation until there are at least 4 protein-ligand pairs in the dataset. This is because PLE optimizes binding pairs in order of binding affinity. Considering the rank correlation with only a few protein-ligand pairs is not particularly useful. This option may be adjusted based on the number of ligands in your particular dataset.

Run command (4) to perform a single simulation and generate a set of PLE models. Each simulation will produce X models, where X is the number of protein-ligand pairs listed in the SAR file. These example output models are in the /outputs/ directory along with a score.sc scorefile.

### Output and analysis

Individual protein-ligand predicted structures are labeled by a protein-ligand pair designation. Wildtype proteins and ligand combinations will be tagged as WT B 1.pdb through WT E 1.pdb where the 1 indicates the docking run it came from. Mutant receptor-ligand pairs will be tagged as 107 A B 1.pdb through 107 A E 1.pdb. The first two parts indicate the mutant residue number and the mutant residue identity respectively. These are followed by the ligand chain designation and the docking run number.

Structures with the same numeric label are based on the same docking simulation and have a common binding pose. The protein interface contacting each ligand are optimized independently. The score.sc file contains all score terms for each simulation across a single row. Generally, individual ligand interface scores are used to rank models, with a negative score indicating a better model. These ligand interface scores are listed as interface delta *, where * corresponds to the protein-ligand prefix tag seen in the PDB files. The values are appended at the end of each output PDB, and also in the scorefile for each protein-ligand pair. One suggestion is for the end user to examine the top ten percent of models for each pair.

## References

1. T. Amemiya, R. Koike, A. Kidera, and M. Ota. PSCDB: A database for protein structural change upon ligand binding. Nucleic Acids Research, 40(D1):554–558, jan 2012.

2. S. DeLuca, K. Khar, and J. Meiler. Fully Flexible Docking of Medium Sized Ligand Libraries with RosettaLigand. Plos One, 10(7):e0132508, 2015.

3. S. R. Ellingson, Y. Miao, J. Baudry, and J. C. Smith. Multi-conformer ensemble docking to difficult protein targets. Journal of Physical Chemistry B, 119(3):1026–1034, 2015.

4. D. Y. Fu and J. Meiler. RosettaLigandEnsemble: A Small-Molecule Ensemble-Driven Docking Approach. ACS Omega, 3(4):3655–3664, 2018.

5. V. Jaakola, M. Griffith, and M. Hanson. The 2.6 angstrom crystal structure of a human A2A adenosine receptor bound to an antagonist. Science, 322(5905):1211–1217, 2008.

6. S. Motta and L. Bonati. Modeling Binding with Large Conformational Changes: Key Points in Ensemble-Docking Approaches. Journal of Chemical Information and Modeling, 57(7):1563–1578, 2017.

7. Z. Yang, S. Han, M. Keller, A. Kaiser, B. J. Bender, M. Bosse, K. Burkert, L. M. Kögler, D. Wifling, G. Bernhardt, N. Plank, T. Littmann, P. Schmidt, C. Yi, B. Li, S. Ye, R. Zhang, B. Xu, D. Larhammar, R. C. Stevens, D. Huster, J. Meiler, Q. Zhao, A. G. Beck-Sickinger, A. Buschauer, and B. Wu. Structural basis of ligand binding modes at the neuropeptide YY1 receptor. Nature, 556(7702):520–524, 2018.

8. A. Zhukov, S. P. Andrews, J. C. Errey, N. Robertson, B. Tehan, J. S. Mason, F. H. Marshall, M. Weir, and M. Congreve. Biophysical mapping of the adenosine A2A receptor. Journal of medicinal chemistry, 54(13):4312–23, jul 2011.

